# Cortical acetylcholine dynamics are predicted by cholinergic axon activity and behavior state

**DOI:** 10.1101/2023.11.14.567116

**Authors:** Erin Neyhart, Na Zhou, Brandon R. Munn, Robert G. Law, Cameron Smith, Zakir H. Mridha, Francisco A. Blanco, Guochuan Li, Yulong Li, Matthew J. McGinley, James M. Shine, Jacob Reimer

## Abstract

Even under spontaneous conditions and in the absence of changing environmental demands, awake animals alternate between increased or decreased periods of alertness. These changes in brain state can occur rapidly, on a timescale of seconds, and neuromodulators such as acetylcholine (ACh) are thought to play an important role in driving these spontaneous state transitions. Here, we perform the first simultaneous imaging of ACh sensors and GCaMP-expressing axons *in vivo*, to examine the spatiotemporal properties of cortical ACh activity and release during spontaneous changes in behavioral state. We observed a high correlation between simultaneously recorded basal forebrain axon activity and neuromodulator sensor fluorescence around periods of locomotion and pupil dilation. Consistent with volume transmission of ACh, increases in axon activity were accompanied by increases in local ACh levels that fell off with the distance from the nearest axon. GRAB-ACh fluorescence could be accurately predicted from axonal activity alone, providing the first validation that neuromodulator axon activity is a reliable proxy for nearby neuromodulator levels. Deconvolution of fluorescence traces allowed us to account for the kinetics of the GRAB-ACh sensor and emphasized the rapid clearance of ACh for smaller transients outside of running periods. Finally, we trained a predictive model of ACh fluctuations from the combination of pupil size and running speed; this model performed better than using either variable alone, and generalized well to unseen data. Overall, these results contribute to a growing understanding of the precise timing and spatial characteristics of cortical ACh during fast brain state transitions.

## INTRODUCTION

While animals are awake, their level of alertness and engagement with the world varies from moment to moment, and these spontaneous state changes can occur on fast timescales that are correlated with behavior and rapidly shift neural activity from one mode to another. For example, in mouse visual cortex, neurons are depolarized and membrane potential variability is reduced during exploratory periods of locomotion and whisking^1–4^, visual responsiveness is enhanced^1,3–7^, and population noise and signal correlation are reduced^8,9^. Outside of these movement periods (i.e. during “quiet wakefulness”) there are also smaller changes in state with similar neural correlates, including a depolarized and less variable membrane potential, reduced amplitude of low-frequency membrane potential oscillations, and increased reliability and selectivity of evoked responses^1,2^. These covert state changes are tracked by dilation and constriction of the pupil that occurs in the absence of any external changes in luminance.

Over the past decade, the mouse has served as an important model for understanding the mechanisms underlying these changes in state, which include neuromodulators^10^ such as acetylcholine (ACh)^11–14^. Indeed, activity in cholinergic axons from the basal forebrain (BF) projecting to the cortex is increased during whisking^15–19^, locomotion^15–17,20–22^, and pupil dilation outside of exploratory periods^20,22^, and genetically-encoded ACh sensors have confirmed some of these effects^21,23–26^. Axonal GCaMP activity might not have a straightforward relationship to ACh levels, which depend on local release probabilities and clearance kinetics, as well as the kinetics of the fluorescent sensors themselves. Thus, more work is required to determine if these measurements are equivalent, and to precisely identify when and where ACh is available to influence the cortex during spontaneous fast state changes. Furthermore, the relationship with pupil and locomotion suggests that it should be possible to build a model that uses these easily measured variables to predict a significant amount of variance in cortical ACh. If this model was shown to generalize well, it could then be employed in other head-fixed mouse experiments to estimate ACh fluctuations in the absence of ground truth data.

We took advantage of recent advances in genetically-encoded fluorescent sensor technology^23^ to address these questions. We performed in-vivo benchmarking of GRAB-ACh 3.0 with exogenous application of ACh, and characterized the relationship with spontaneous running speed and pupil size fluctuations. We performed the first simultaneous measurements of axons and ACh activity. Consistent with expectations that cholinergic axons indeed releasing ACh, we found that ACh and axonal activity are tightly correlated, and that on a fine spatial scale larger ACh increases can be observed closer to cholinergic axons compared to further away. We demonstrate an approach for deconvolving GRAB-ACh fluorescence to estimate the underlying changes in ACh levels – an approach which we believe could also be usefully applied to other fluorescent sensors that provide measurements of “analog” signals like neuromodulator or neuropeptide levels. Finally, we show that ACh levels can be predicted from pupil size and treadmill speed with relatively high accuracy by a trained neural network.

## RESULTS

### Experimental setup and in vivo sensor validation

We used two-photon microscopy to measure ACh activity in the cortex of wild type mice by implanting a cranial window over the visual cortex (VIS) and expressing the genetically encoded fluorescent ACh sensor GRAB-ACh3.0 via viral injection (Fig. 1A). Although GRAB-ACh3.0 has been benchmarked against varying concentrations of ACh *in vitro*^23^, this benchmarking has not been performed *in vivo*. To measure the sensor response *in vivo,* we utilized an approach similar to two-photon-guided patching^1^ with a pipette entering through a hole in the coverslip. This technique allowed us to image GRAB-ACh signals in an anesthetized mouse while directly puffing small amounts of ACh into the cortex in the field of view to measure ACh sensor responses (Fig. 1B, Supplemental Video 1). We observed strong increases in GRAB-ACh fluorescence with 10µM, 100µM, and 1mM ACh concentrations (Fig. 1C). GRAB-ACh ΔF/F responses appeared to be larger with puffs of 100µM ACh than 1mM ACh, but this might be due to a saturating response of the sensor at 1mM and the higher ACh concentration not fully washing out before the next puff was administered. Puffs of ACh-free cortex buffer and 100µM norepinephrine resulted in mild decreases in GRAB-ACh fluorescence, presumably due to the transient wash-out of endogenous levels of ACh. Interestingly, puffs of 1µM ACh also resulted in a decrease in fluorescence, which might indicate that baseline concentrations of ACh under isoflurane anesthesia in visual cortex are above 1µM. Overall, these experiments directly confirmed that changing ACh levels in vivo are rapidly tracked by the GRAB-ACh sensor.

**Figure 1:**
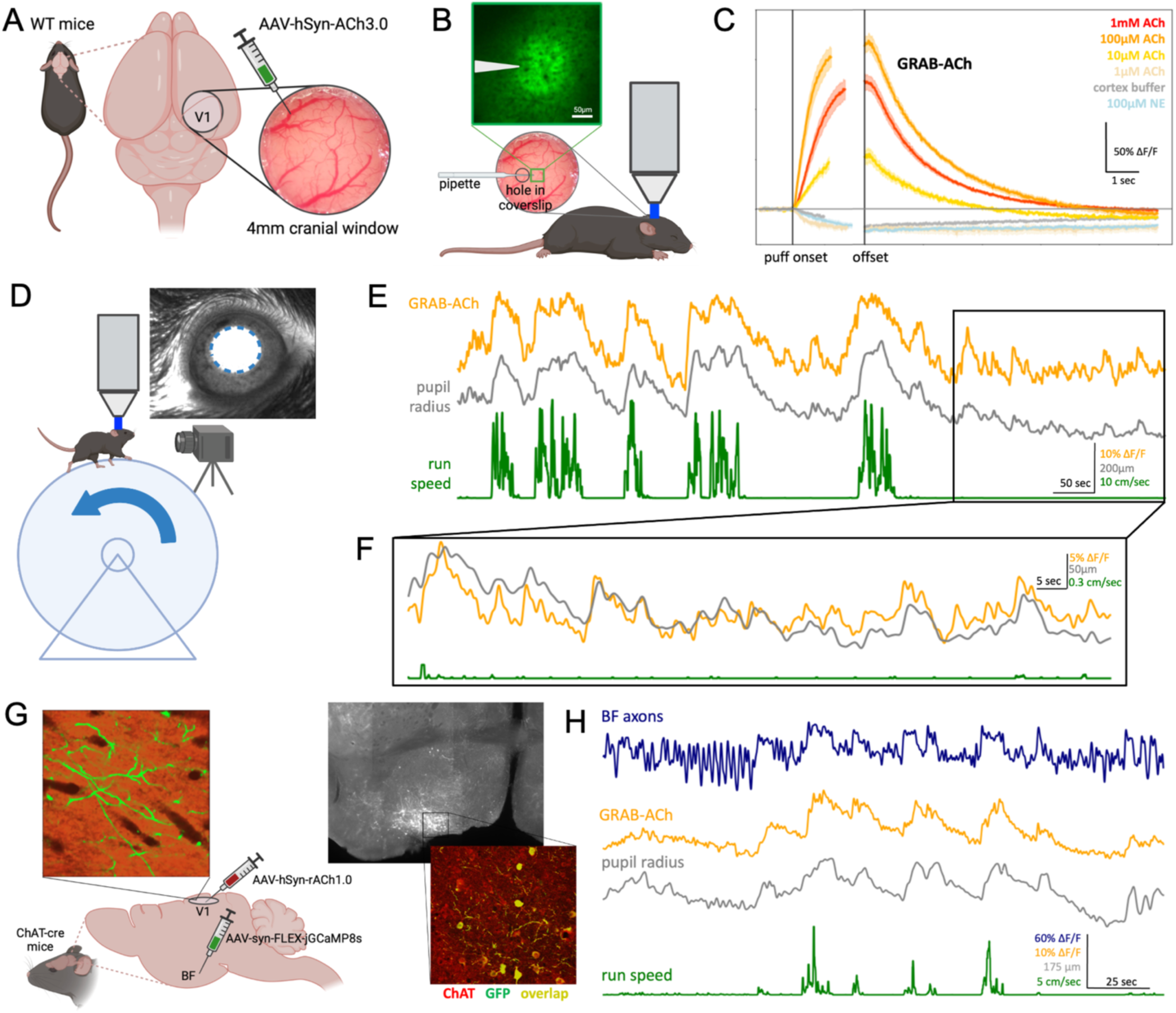
GRAB-ACh activity reveals spontaneous fluctuations in ACh levels which track behavior state and cholinergic axon activity. **A)** Animal prep: a cranial window was implanted over the visual cortex of WT mice and GRAB-ACh3.0 was expressed via viral injection. **B)** For pharmacological validation experiments, a coverslip with a hole was implanted, through which a glass pipette could be inserted. Various concentrations of ACh were puffed onto the cortex while imaging was performed under isoflurane anesthesia. **C)** Puffs of varying concentrations of ACh resulted in increases in GRAB-ACh fluorescence, while injections of control substances (cortex buffer, NE) resulted in decreases in fluorescence. Puffs of 1µM ACh also resulted in a decrease in fluorescence, consistent with an ambient ACh concentration above 1µM. (N puffs in each group in the order listed = 10, 11, 5, 18, 15, 5; error bars represent standard error). **D)** Experimental setup for awake, behaving experiments. The mouse was head-fixed on a treadmill in a dark room underneath a two-photon microscope while treadmill speed and a video of the eye were recorded. **E)** Example traces of spontaneous ACh levels, pupil radius, and run speed while the mouse naturally transitioned between states. Run periods were accompanied by large increases in pupil size and ACh level (as indicated by GRAB-ACh fluorescence). **F)** A close-up of the example trace in E), highlighting the close correspondence of ACh with spontaneous pupil fluctuations during a period of stillness (quiescence). **G)** Animal prep for axon imaging experiments. GCaMP8s was expressed in cholinergic axons by viral injection into the horizontal limb of the diagonal band (HDB) in the basal forebrain (BF) of ChAT-cre mice. A cranial window was implanted over the visual cortex and rACh was expressed via viral injection. Verification of injection sites was subsequently done histologically. **H)** Example traces of spontaneous axon activity in comparison with simultaneous ACh, pupil radius, and run speed, in which correspondence between all four traces is apparent. Panels A, B, D, and G created with BioRender.com

We next repeated the measurements we had previously performed relating cholinergic axons to locomotion and pupil under spontaneous conditions, but now using the GRAB-ACh sensor. The mouse was head-fixed on a treadmill and allowed to run freely while the pupil was recorded with a camera, and we simultaneously imaged GRAB-ACh in VIS (Fig. 1D). As expected based on previous studies^15–17,20–22^, we observed strong increases in ACh levels which tracked spontaneous locomotion periods and the accompanying large pupil dilations (Fig. 1E). Focusing in on the small dilations that occurred outside of locomotion (i.e., during quiet wakefulness), we observed a high correlation between spontaneous ACh fluctuations and small dilations and constriction of the pupil (Fig. 1F). This was surprising, as previous work^27^has emphasized the relationship between pupil and norepinephrine, and we did not expect there to be such a close correspondence between pupil and cortical ACh measured with GRAB-ACh.

This raised the question of whether we could precisely define the temporal relationship between changes in ACh levels and cholinergic axon activity by measuring them simultaneously. While neuromodulator axon activity and release have been measured simultaneously in vitro^28^, to our knowledge this experiment has not been performed in vivo for any neuromodulator. We expressed FLEX-GCaMP8s in cholinergic axons via viral injection into the horizontal diagonal band (HDB) of the BF in ChAT-cre mice, as well as the red ACh sensor rACh1.0 in VIS (Fig. 1G). HDB was chosen due to its preferential projections to VIS^24,29,30^, and we used rACh1.0 so that the sensor could be simultaneously imaged with GCaMP activity on red and green emission channels. We observed that BF axon activity was more transient and phasic than ACh activity, but both closely tracked locomotion periods and pupil size (Fig. 1H). Thus, cholinergic axon activity is indeed tightly related to local ACh levels in the cortex, validating previous studies that have used axon activity alone to measure cholinergic tone^16,20^ and ameliorating concerns about mechanisms that could potentially dissociate axonal action potentials from neuromodulator release^31^.

### Changes in ACh levels surrounding periods of locomotion

As expected from previous studies^21,23,25,26^, ACh levels in VIS increased during running. To more closely examine the timing, magnitude, and spatial heterogeneity of this increase, we measured and compared ACh activity during spontaneous running across three cortical areas: VIS, somatosensory cortex (SS), and motor cortex (MO). When aligning ACh activity to all run onsets and offsets, we observed a similar pattern across all three areas: a large increase right before run onset and a slower decline back to baseline after run offset (Fig. 2A). To look at the timing of these onsets and offsets more closely, we restricted to isolated running periods with at least 15 seconds of quiet wakefulness before onsets or after offsets. Because recordings in VIS, SS, and MO were not conducted simultaneously, we matched run periods by similar speed. We found that ACh activity began to increase 1.37 +/- 0.4 seconds before run onset and declined after running with a decay constant of 7.2 +/- 1.4 seconds, and this timing was similar across all three brain areas (Fig. 2B). The timing of the initial ACh rise (within the first 5 seconds of running) was not dependent on run speed (Fig. 2C). In contrast with a previous study^21^, we observed a clear relationship between run speed and duration and the magnitude of ACh changes, with faster and longer run periods leading to larger ACh increases than slower and shorter run periods (Fig. 2D). Run speed, duration, and magnitude of ACh response in the final 5 seconds of running did not have a significant effect on decay time (Fig. 2E). Again, to compare ACh response magnitude across cortical areas, we matched run periods by run speed and duration (Fig. 2F). While the maximum ΔF/F response was significantly different across brain areas when binning by run speed and comparing medians (Kruskal-Wallis test, p=0.0065), there was no difference when comparing means (one-way ANOVA, p=0.188). When binning by run duration, there was no significant difference across areas when comparing either medians (p=0.719) or means (p=0.811). In summary, we observed the expected rapid increase of ACh preceding running and a slower decrease back to baseline right at run offset; the magnitude of this increase varied by run speed and duration but was relatively homogeneous across cortical areas.

**Figure 2:**
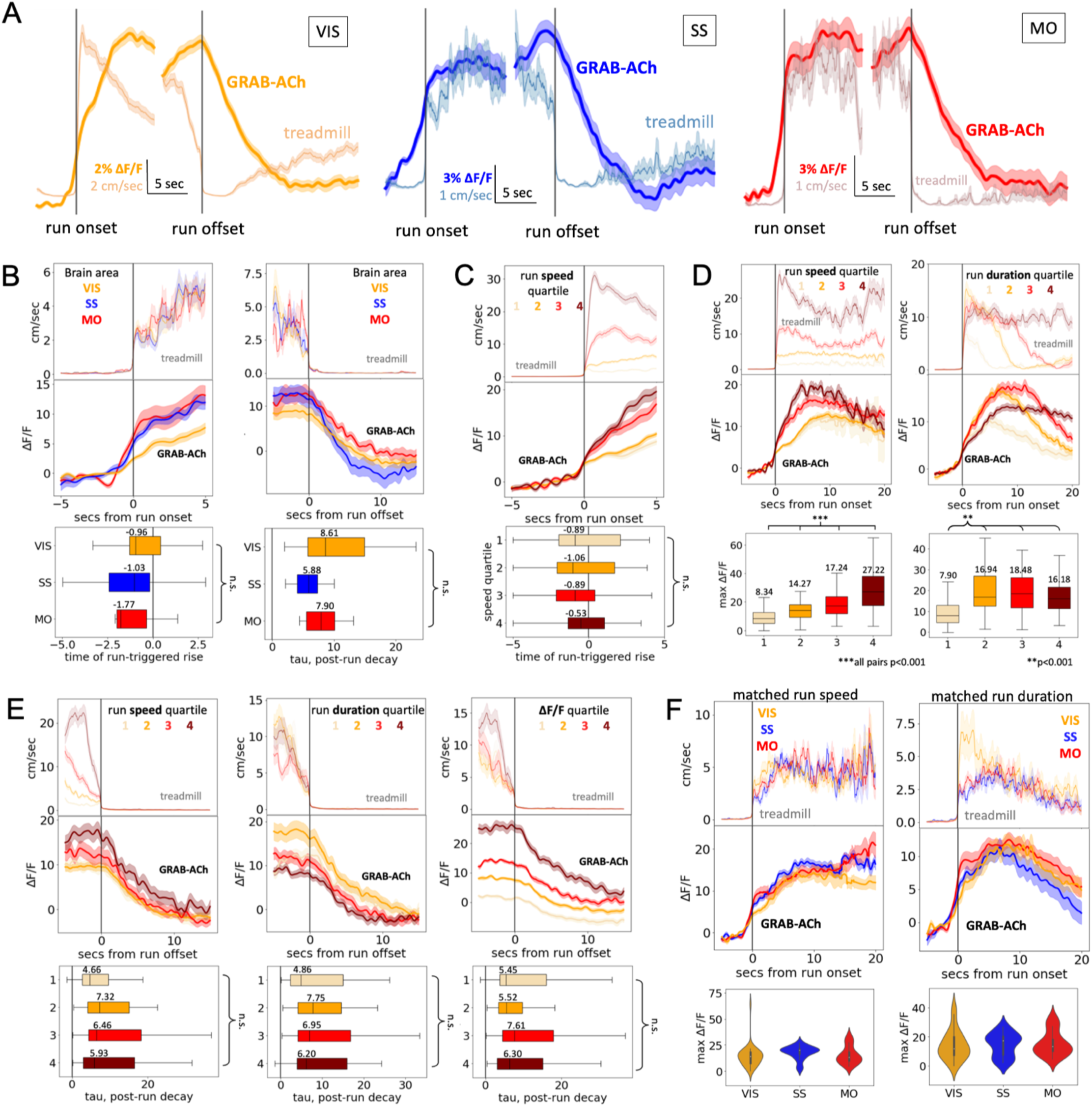
Locomotion is linked to large-scale, slowly decaying increases in ACh which vary by run magnitude and are relatively homogeneous across the dorsal cortex. **A)** Mean run onset- and offset-triggered ACh traces for VIS (n=694 run periods), SS (n=70), and MO (n=51). **B)** Mean run onset- (n=24 in each area) and offset- (n=14) triggered ACh in VIS, SS, and MO (matched by mean run speed in the first/last 5 seconds of running). Differences in median rise time and median decay tau were not significant across areas (overall onset p=0.375, offset p=0.126, Kruskal-Wallis test). **C)** In VIS, mean run onset-triggered ACh binned by run speed in the first 5 seconds of running (n=80-81 in each bin; see Supp. Fig. 1 for a breakdown of run speeds). The median time of ACh rise in response to running was not significantly different across groups (p=0.629). **D)** Mean run onset-triggered ACh in VIS binned by mean speed in entire run period (n=100-101 in each in each bin) or total run duration (n=99-100). In both conditions, the median peak ACh ΔF/F during running was significantly different across bins (overall p<0.001; binning by run speed, all pairs p<0.001; binning by run duration, 1 vs. 2, 1 vs. 3, & 1 vs. 4 p<0.001; all other pairs n.s.) **E)** Mean run offset-triggered ACh in VIS binned by mean run speed in last 5 seconds of running, by total run duration, or by mean ΔF/F in last 5 seconds of running (n=50-51 in each bin). Median decay tau of ACh (fitted to first 10 seconds after run offset) was not significantly different across bins in all three conditions (left-to-right overall p=0.181, 0.478, 0.475). **F)** Mean run onset-triggered ACh in each brain area matched by mean speed in entire run period or total run duration (n=37 in each area), and violin plots showing the medians and distributions.

### Changes in ACh levels surrounding dilation events during quiescence

As we have done previously^1,20^, we next excluded active periods to focus on the smaller fluctuations in pupil size that occurred outside of running. Some previous studies have found that cholinergic axon activity^20,22^ or ACh levels^24^ are coherent with and precede changes in pupil size in cortex, while others found that pupil size was in general not a good predictor of cortical ACh activity^16,25^. In our hands, we found that ACh fluctuations during quiet wakefulness were tightly tracked by small pupil dilations (Fig. 3A). ACh activity was highly coherent with pupil size across brain regions, both during quiescence (Fig. 3B) and when including periods of locomotion (Supp. Fig. 2A). When we compared dilation periods of similar magnitude (Fig. 3C), we found that larger dilation periods were accompanied by larger ACh increases, as well as a shorter lag between ACh rise and dilation onset. Comparing dilation periods of similar magnitude across cortical areas (Fig. 3D), our results suggest that dilation-linked ACh increases had a shorter lag in VIS, and were larger in SS. Thus, cortical ACh levels are correlated with and precede pupil dilation, although the timing and magnitude of this relationship may be somewhat heterogeneous across areas.

**Figure 3:**
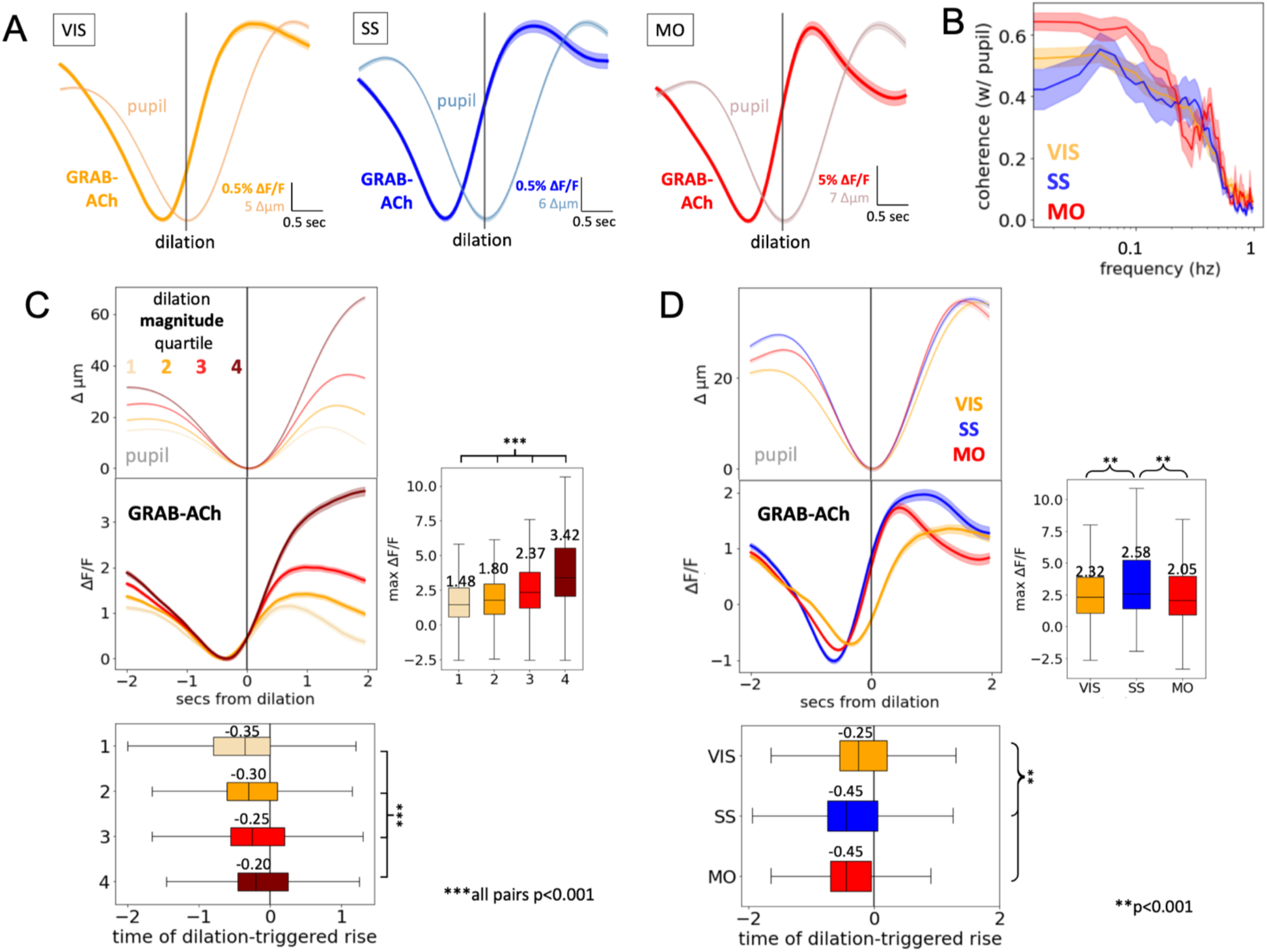
Pupil dilation during quiescence is linked to small-scale, rapid changes in ACh which vary by cortical area. **A)** Dilation-triggered ACh traces during stillness for VIS (n=6254 dilations), SS (n=1644), and MO (n=1539). **B)** Mean coherence between pupil radius and ACh in different brain areas during quiescence. Across areas, coherences exhibited a broad peak at lower frequencies (<0.1 hz). **C)** In VIS, dilation-triggered ACh traces sorted by dilation magnitude (n=1563-1564 in each bin). Median time of dilation-triggered ACh rise was significantly different across bins (overall p<0.001, all pairwise comparisons p<0.001, Kruskal-Wallis test). Median peak ACh ΔF/F was also significantly different across bins (overall p<0.001, all pairwise comparisons p<0.001). **D)** Dilation-triggered ACh traces in VIS, SS, and MO matched by dilation magnitude (n=1539 in each area). Median time of dilation-triggered ACh rise was significantly closer to dilation start in VIS (overall p<0.001, VIS vs. SS and VIS vs. MO p<0.001, SS vs. MO p=0.072, Kruskal-Wallis test). Median peak ACh ΔF/F was also significantly higher in SS than in VIS or MO (overall p<0.001, SS vs. VIS and SS vs. MO p<0.001, VIS vs. MO p 0.164).

### Temporal relationship between ACh and cholinergic axons

Previous studies have used calcium indicators expressed in axons as a proxy for changes in local ACh levels^15,16,20,22^. However, axon activity may be dissociated from extracellular neuromodulator levels, which depend on release probabilities and clearance kinetics. In order to directly measure the relationship between axon activity and fluctuations in ACh levels, we recorded cholinergic axon activity with GCaMP in the green channel and rACh in the red channel. As we have observed previously^20^, axon activity increased around running periods. Consistent with the hypothesis that changes in ACh levels are driven by local cholinergic axon activity, we observed that changes in axon activity preceded changes in ACh activity both at the onset and offset of running (Fig. 4A). When we quantified this timing by examining run periods with no running in the 15 seconds before or after running, we found that axon activity began increasing 1.03 seconds before the rise in ACh, and GRAB-ACh fluorescence decayed 4.4 times as slow as the GCaMP axon activity after running, despite the similar off-kinetics of GCaMP and the rACh sensor (Fig. 4B). Although not unexpected given previous observations of axons and sensors in other experiments, this is the first direct evidence that cholinergic GCaMP activity indeed is correlated with local ACh release, and that ACh remains available for several seconds after axonal activity returns to baseline for large ACh transients.

**Figure 4:**
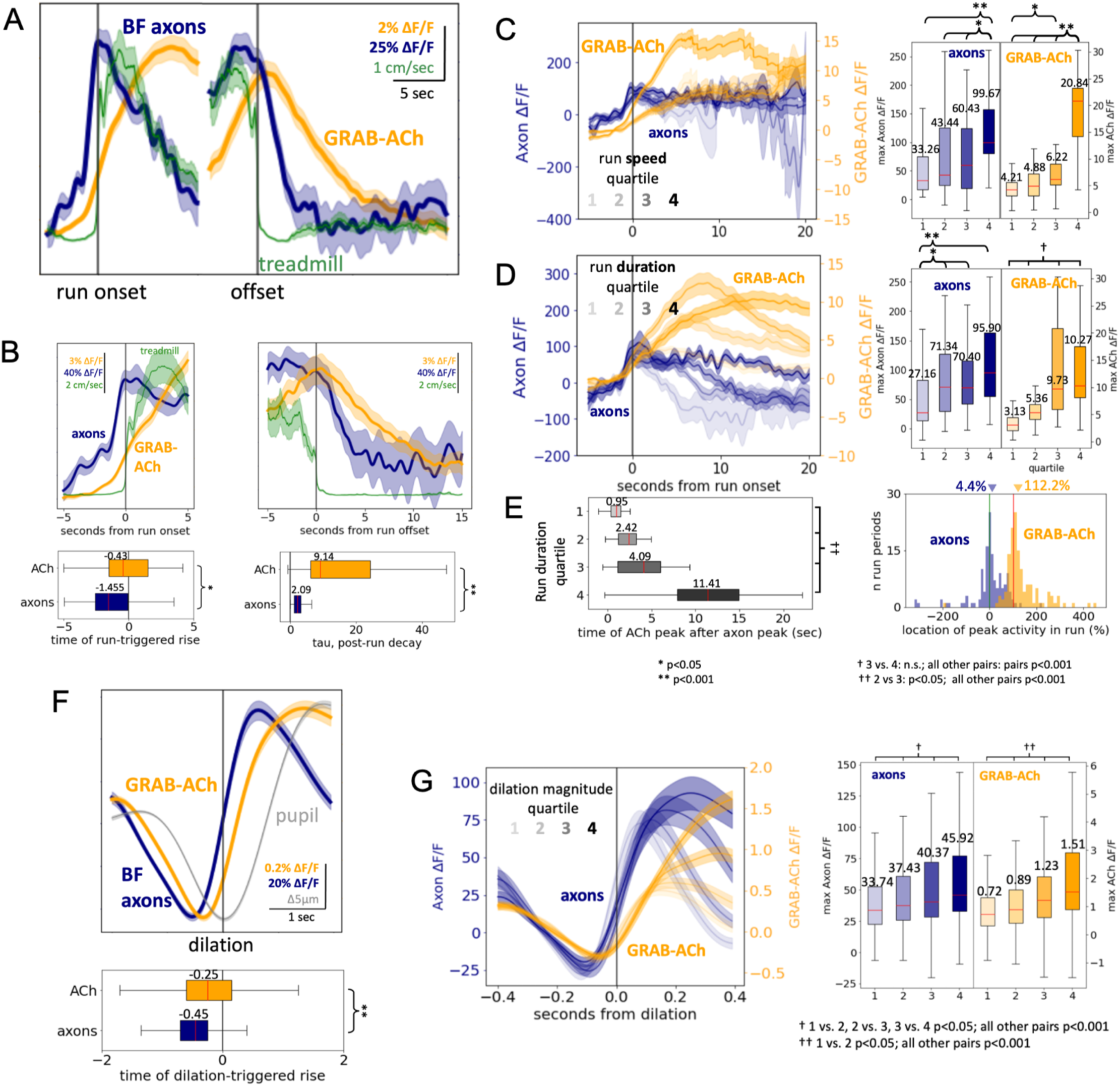
Cholinergic axon activity is accompanied by increases in local ACh levels which precede locomotion onset and pupil dilation. **A)** Mean run onset- and offset-triggered ACh and axon traces (n=242 run periods). **B)** Mean run onset- (n=94) and offset- (n=45) triggered ACh and axon activity for run periods with no running 15 seconds before or after running were also excluded. Median time of increase in ACh activity was significantly later than time of increase in axon activity (p=0.011, Kruskal-Wallis test). Median decay tau was also significantly faster for axon activity than ACh activity (p<0.001). **C)** Mean run onset-triggered ACh and axon activity binned by mean run speed (n=39-40). There was a significant difference in median peak ΔF/F across bins for both axon and ACh activity (overall p<0.001 for both groups; see figure for pairwise comparisons.) **D)** Mean run onset-triggered ACh and axon activity binned by run duration (n=39). There was a significant difference in peak ΔF/F across bins for both axon and ACh activity (overall axons p=0.001; ACh p<0.001; see figure for individual pairwise comparisons). **E)** Left: Time of peak ACh ΔF/F after peak axon ΔF/F for the run bins in D). There was a significant difference across bins (overall p<0.001, 2 vs. 3 p=0.42, all other pairs p<0.001.) Right: histogram of location of peak axon vs. peak ACh peri-run ΔF/F, expressed as a percentage of run (where 0% is peak occurring at run onset and 100% is peak occurring at run offset). Axon traces peaked early, at 4.4% of the run (median across axons), whereas GRAB-ACh signals peaked later, at 112.2% of the run. **F)** Dilation-triggered ACh and axon traces during stillness (n=1678 dilation periods). Median time of increase in ACh activity was significantly later than time of increase in axon activity (p<0.001). **G)** Dilation-triggered ACh and axon traces binned by dilation magnitude (n=419-420 in each bin). There was a significant difference in median peak ΔF/F for both axon activity (overall p<0.001, 2 vs. 3 p=0.054, all other pairs p<0.001) and ACh activity (overall p<0.001, 1 vs 2 p=0.005, all other pairs p<0.001).

This led to the question of whether there was an effect of running magnitude on the accompanying axon or ACh activity. Previous evidence for this has been mixed, with one study suggesting that ACh levels do vary by run speed^23^ and another study suggesting neither ACh nor axon activity vary by run speed^21^. When binning by average run speed we found that faster runs resulted in a larger increase in axon activity as well as ACh measured with the fluorescent sensor (Fig 4C). We found a similar pattern when binning by run duration: longer runs resulted in higher magnitude axon and GRAB-ACh activity (Fig. 4D). Interestingly, we noticed that the relative timing of these peaks in run-triggered axon and ACh activity was not consistent across runs of different durations. For longer run periods, the peak in ACh activity occurred significantly later than the peak in axon activity for that same run period (Fig. 4E). Whereas axon activity typically peaked in about the first 4% of the run period, ACh activity did not reach its peak until just after the very end of the run period. Overall, these results suggest that ACh levels may continue to increase under steady-state axonal release, potentially due to saturation of clearance mechanisms.

Next, we examined simultaneously recorded axon and sensor dynamics with respect to pupil fluctuations during periods of quiescence. With respect to dilation onset, axon activity began increasing about 200 milliseconds before ACh sensor activity (Fig. 4F), and larger dilations resulted in a significantly larger magnitude of both axon and ACh sensor fluorescence (Fig. 4G). This suggests that these smaller ACh changes associated with pupil in the absence of running likely occur in a range where ACh is cleared rapidly, and thus tracks more closely with axonal activity. These smaller changes in ACh had a higher cross-correlation with pupil size than did axon activity (Supp. Fig 2D & E), indicating that spontaneous fluctuations in pupil size more closely track ACh changes than cholinergic axon activity. Furthermore, while rACh exhibited peak coherence with pupil size at lower frequencies (<0.1 hz) both during quiescence and when accounting for running periods, cholinergic axon activity was most coherent with pupil size between 0.1-1hz (Supp. Fig. 2B & C).

### Spatial relationship between ACh and cholinergic axons, and temporal modelling

We wondered if our imaging configuration would provide sufficient resolution to observe spatial features of local ACh release surrounding cholinergic axons, which might be difficult given that we only monitored the activity of axons in a 2D plane (i.e. we could potentially be missing axons above and below the field of view; see Supplemental Video 2). In each scan, we binned pixels according to distance from the nearest axon in the field of view (example scan, Fig. 5). Indeed, when looking at spike-triggered ACh in each of these spatial bins, weaker responses were observed in spatial bins further away from the nearest axon (Fig. 5B). The order of response magnitude from nearest to furthest bins was significant (Spearman’s rank correlation coefficient = -0.066, p<0.001). This suggests that there is a small amount of spatial heterogeneity of ACh activity around cholinergic axons, consistent with a volume transmission mechanism.

**Figure 5:**
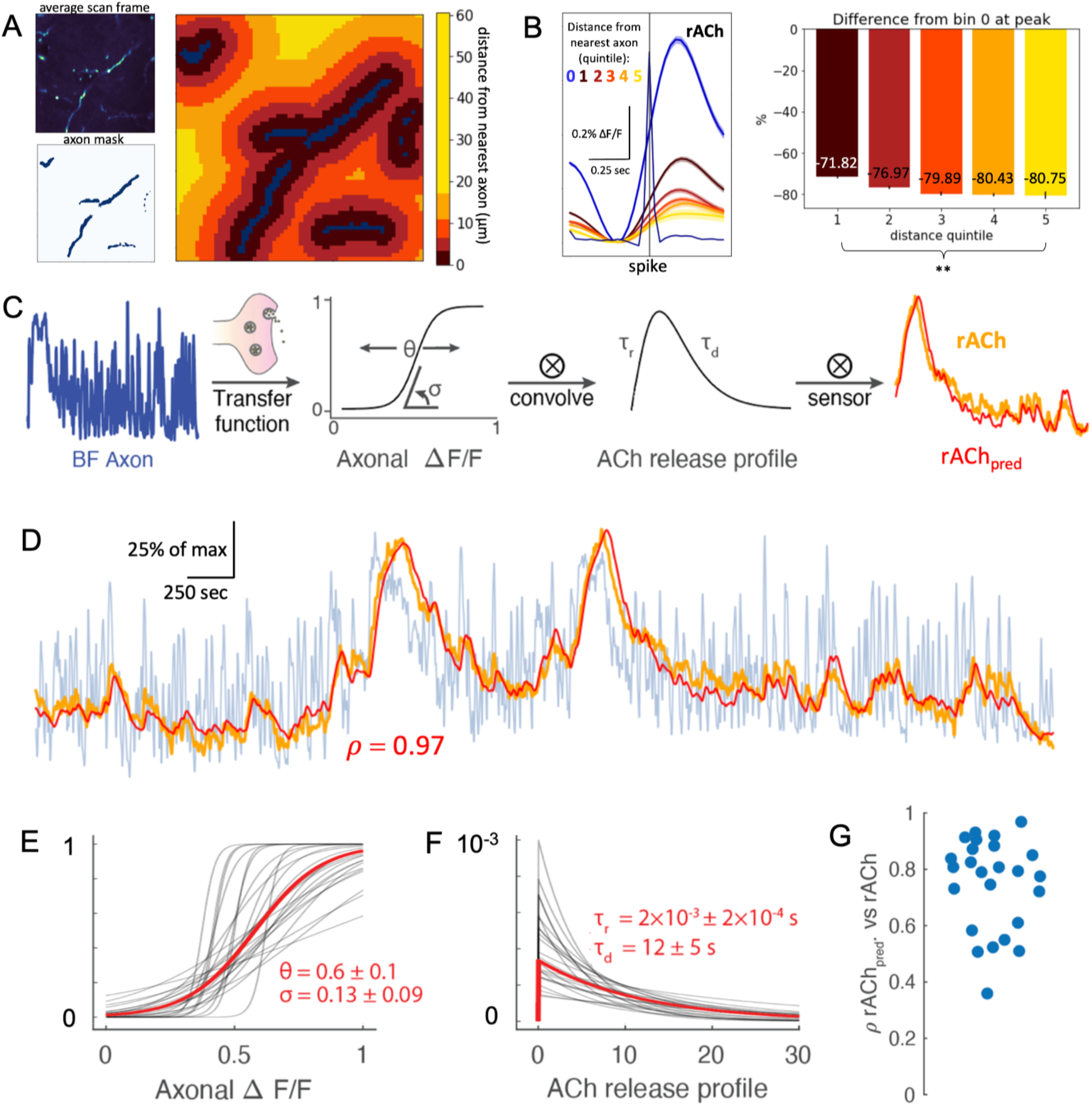
ACh activity decreases at distances further from an axon and can be predicted temporally from cholinergic axon activity. **A)** For one example scan, average scan frame, ROI mask representing location of axons, and a map representing pixel-wise distance from nearest axon as defined by the axon mask. **B)** Left: Mean spike-triggered ACh ΔF/F for all pixels in all scans (n=6747 spikes, ISI = +/- 0.5 sec) binned by distance from nearest axon, where bin 0 represents pixels containing an axon mask. Right: For each bin, mean ΔF/F value at peak activity in bin 0 expressed as percent change from bin 0. The rank order was significant (Spearman rank order correlation coefficient = -0.066, p<0.001). **C)** Prediction model workflow going from recorded cholinergic axon activity to predicted rACh signal. **D)** An example segment of recorded axon activity (blue), recorded rACh activity (orange), and predicted rACh activity (red). Pearson correlation *ρ* = 0.97 between recorded and predicted rACh. **E)** Sigmoidal activation function from axon activity to ACh release for each recording (black) and averaged over all recordings (red). **F)** ACh release functions for each recording (black) and averaged over all recordings (red). **G)** Pearson correlation between empirical and predicted rACh for each recording.

In order to quantify the extent to which GRAB-ACh fluorescence fluctuations could be predicted from axon activity, we built a simple model with a sigmoidal function relating GCaMP axonal activity to ACh release, and an exponential rise (*τ_r_*) and decay ^32^(*τ_d_*; Fig. 5C) for ACh. We found that this simple dynamical model was able to use axonal fluorescence (Fig. 5D, blue) to generate a prediction for rACh (Fig. 5D, red) that closely resembled empirically measured rACh (Fig. 5D orange; Pearson correlation *ρ* = 0.97 between observed and predicted rACh). Overall, we found the optimization algorithm predicted a slightly right-shifted sigmoidal (i.e., more calcium required for a response; Fig. 5E) and a rapid release (*τ_r_*∼0) with decay over seconds (*τ_r_*∼12s, Fig. 5F). Across all recordings the algorithm could accurately predict rACh fluorescence (*ρ̅* = 0.8 ± 0. 2; Fig. 5G) and explain a significant amount of the rACh variance (*ρ̅*^#^ = 0.6 ± 0. 2). This algorithm provides a simple method to use recordings of axonal fluorescence to predict what the GRAB-ACh sensor would report. More generally, it provides a strong validation for existing and future studies using cholinergic axon activity as a proxy for measuring local ACh levels in cortex, with the caveat that ACh levels and axon activity levels may not be linearly related for large cholinergic events.

### Deconvolution of GRAB-ACh sensors

Having confirmed that cortical ACh levels track state-related variables and cholinergic axon activity, we wanted to better understand the actual time course of ACh fluctuations independent of GRAB-ACh sensor kinetics. We estimated the time course of [ACh] by deconvolving the GRAB-ACh impulse response measured in vitro from the fluorescence trace (Fig. 6A; more details in Methods). Applying this deconvolution approach to ACh puffs (as in Figure 1) yielded the expected faster rise and return-to-baseline of [ACh] than the GRAB-ACh sensor (Fig. 6B). Large-scale ACh changes such as those surrounding running were too slow to be substantially affected by the sensor off time (Fig. 6C & D). However, deconvolution emphasized that the rapid clearance of smaller changes in ACh levels (such as those associated with pupil dilation in the absence of running) may be obscured by the off kinetics of the GRAB-ACh sensor (Fig. 6E-H). By chance, the sensor kinetics for GCaMP and rACh were similar enough that the lag between them was similar for e-convolved and non-deconvolved traces (Fig. 6G).

**Figure 6:**
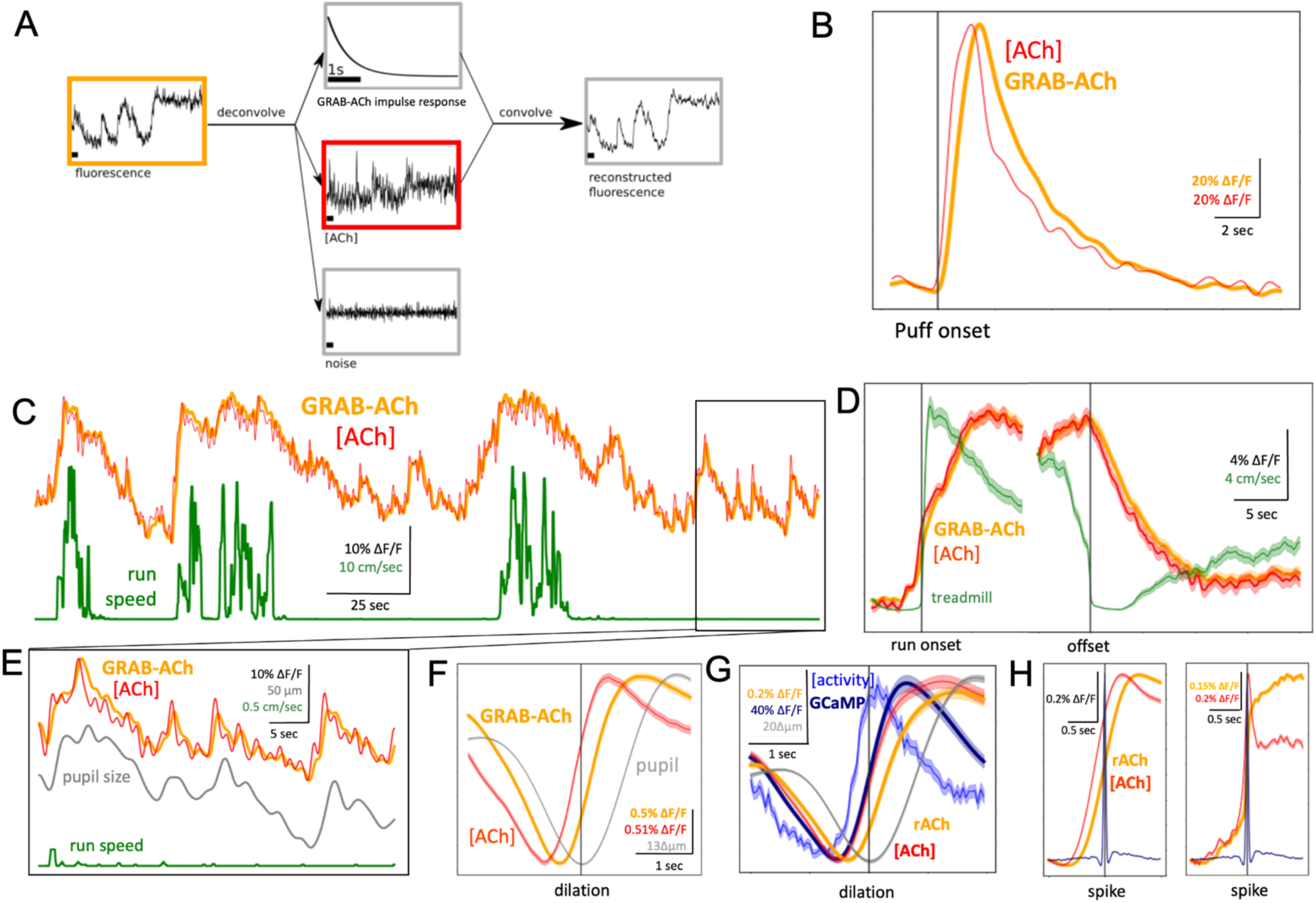
For small and fast events, GRAB-ACh deconvolution can help reveal rapid ACh kinetics. **A)** Schematic illustrating the causal deconvolution filter. **B)** A single 100µM ACh injection-triggered GRAB-ACh trace with the deconvolved ACh trace overlaid. As expected, deconvolved ACh reaches a peak slightly before GRAB-ACh ΔF/F. **C)** An example trace of GRAB-ACh in VIS, along with deconvolved ACh and running trace. Deconvolution does not substantially alter the large-scale changes of ACh seen around running. **D)** Run onset- and offset-triggered GRAB-ACh and deconvolved ACh for all running periods in VIS recordings. Deconvolution does not substantially alter the timing of ACh increases before running or decay after running. **E)** Close-up of a quiescence period in the example traces in C), with the addition of pupil size. Deconvolution reveals sharper, more phasic activity of ACh correlated with spontaneous pupil size fluctuations. **F)** Dilation-triggered GRAB-ACh and deconvolved ACh traces for all dilation periods in VIS recordings. Deconvolved ACh rises and peaks sooner than is indicated by GRAB-ACh fluorescence. **G)** Dilation-triggered rACh and axon activity (GCaMP) with deconvolved ACh and axon activity traces. After deconvolution, the order of axon activity and ACh activity revealed in previous experiments is preserved. **H)** Spike-triggered GRAB-ACh and deconvolved ACh for all simultaneous axon-GRAB-ACh recordings (n=45,669 spikes; left, 1hz lowpass filtered; right, unfiltered). Again, deconvolved ACh rises and peaks sooner than what is indicated by GRAB-ACh fluorescence.

### Prediction of GRAB-ACh activity from locomotion and pupil size

Finally, given our results indicating a close relationship between ACh and state-related behavior variables, we wondered how accurately we could predict ACh levels from these variables. To this end, we trained a long short-term memory (LSTM) recurrent neural network to predict GRAB-ACh fluctuations from the recorded behavioral dynamics – pupil size and running speed and their temporal derivative (Fig. 7A). The statistical model was able to reproduce the GRAB-ACh from training recordings (*ρ̅* = 0.72 ± 0. 21; Fig. 7B), explaining over 50% of the GRAB-ACh variance (*ρ̅*^2^ = 0.57 ± 0. 26), both during high-arousal states marked by locomotion as well as quiescent states (Fig. 7A right). As expected, the model’s explanatory capacity improved with an increase in correlation between the behavioral variables and GRAB-ACh signal, with model accuracy routinely exceeding behavioral signal by itself (Fig. 7C). Importantly, the model was able to correctly predict GRAB-ACh from behavior in unseen data (held-out validation dataset) at a performance matching training data (Fig. 7D left; explained-variance *ρ̅*^2^ = 0.60 ± 0. 23), indicating that it is general enough to be useful for predicting ACh levels from recordings without ground truth training data, i.e. in any head-fixed mouse experiment with recordings of treadmill and pupil size. As both behavioral variables are used as proxies for arousal, we wanted to contrast their explanatory power. Using either behavioral metric (pupil or locomotion speed) independently led to a decrease in signal accuracy compared to using both variables (Fig. 7D right pupil p = 3 × 10^$%^, speed p = 8 × 10^$&^ paired t-test). Overall, we have demonstrated that behavioral variables used to demarcate changes in brain state can also be used to predict ACh levels in the cortex.

**Figure 7:**
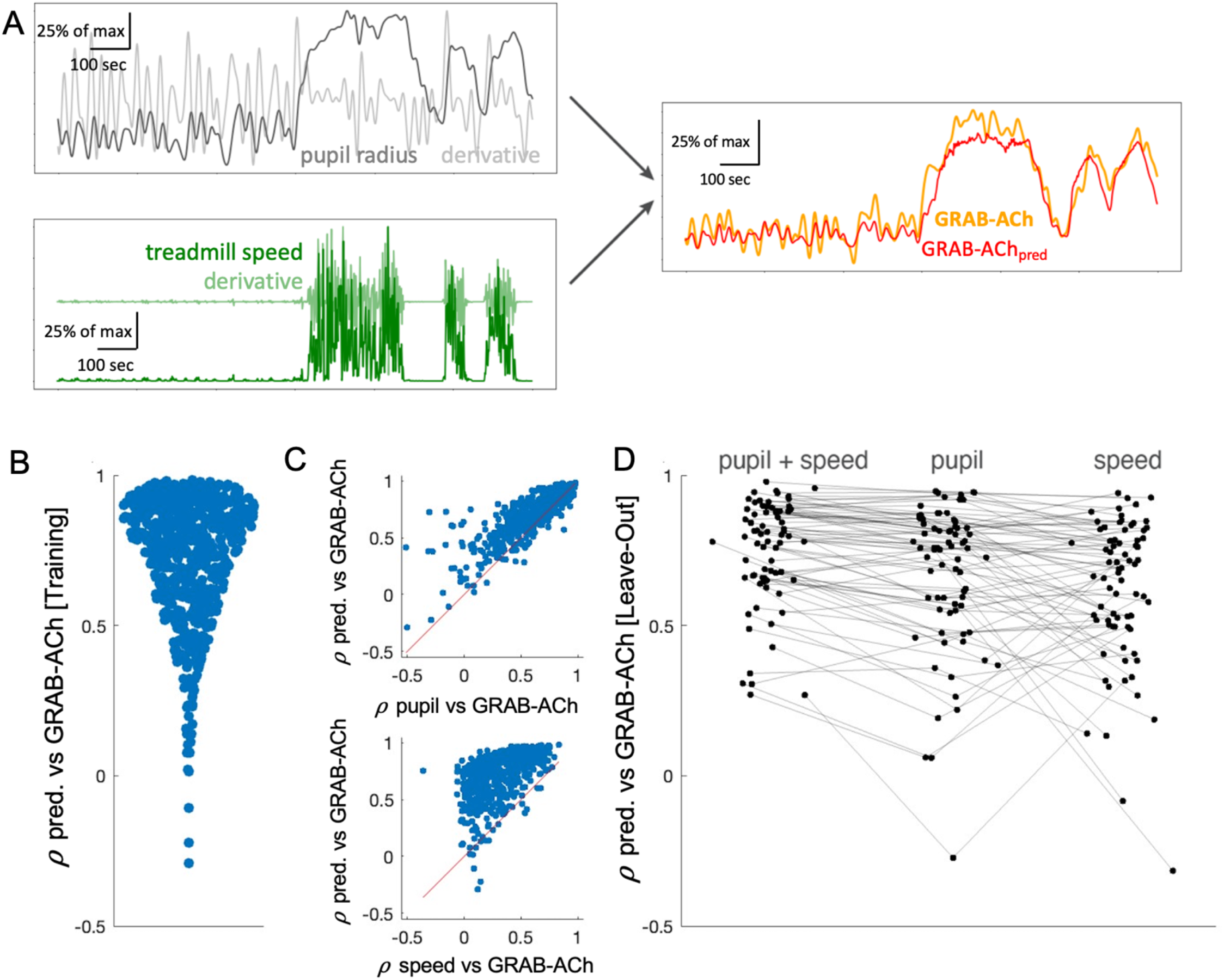
ACh levels can be predicted from pupil size and locomotion speed. **A)** A model was constructed from pupil and treadmill measurements and their derivatives. Example traces of pupil radius/derivative and treadmill speed/derivative and the accompanying observed GRAB-ACh activity with predicted GRAB-ACh. **B)** Correlation between observed and predicted GRAB-ACh for each recording (training data). **C)** (top) For each recording, correlation between pupil size and observed GRAB-ACh versus correlation between predicted and observed GRAB-ACh (76.9% above the diagonal); (bottom) the same comparison, but instead with treadmill speed (97.5% above the diagonal). **D)** For the leave-out data, correlations between observed and predicted GRAB-ACh when using as prediction variables pupil size and treadmill speed together, or either of these two variables alone.

## DISCUSSION

Our study leverages recent advances in genetically encoded fluorescent indicators to investigate the question of when and where ACh is available during spontaneous, fast brain state transitions. Consistent with previous work^21,23,25,26^, we found that transitions from quiet wakefulness to the high-arousal state during locomotion resulted in a large increase in ACh which outlasted running periods by several seconds. Also consistent with previous results^24^, small ACh increases occurred around a half second before the onset of spontaneous epochs of pupil dilation that occur when animals are sitting quietly.

Previous studies have presented mixed results regarding whether or not there is positive association between run speed and ACh response^21,23^. Here, we found that faster and longer run periods do indeed result in a larger ACh response. Differences in findings between our and previous studies could be due to the exact time point where ACh is measured relative to running speed, since we find that the peak in ACh response occurs late in the running period. Similarly, we found a clear positive association between pupil dilation magnitude and ACh response in our data, a point on which some studies differ^24,25^.

We sought to examine the heterogeneity of ACh release and clearance across several areas of the dorsal cortex. While the BF is often referred to as a monolithic structure, evidence suggests it is actually somewhat modular with segregated projections to the cortex^33,34^. VIS receives a large proportion of its cholinergic projections from HDB, while the posterior nucleus basalis (NB) preferentially targets the auditory and anterior NB targets the SS^29^. A recent study^25^ found heterogeneity in ACh responses across the dorsal cortex, specifically for run-triggered ACh. In contrast, we found that the magnitude and timing of large-scale run-evoked ACh was largely homogeneous across the dorsal cortical areas we examined; this contrasting result may be due to differences in the way we computed these metrics or due to differences in experimental setup (e.g., one- vs. two-photon or differences in running measurements). On the other hand, we found that small-scale ACh increases – linked to pupil dilation – were larger in visual cortex than in the more anterior cortical areas, and occurred with less of a delay relative to dilation.

To assess whether there was a direct relationship between ACh levels and cholinergic axon activity, we performed simultaneous measurements of cholinergic axons using GCaMP and the accompanying changes in ACh levels using a fluorescent ACh sensor. To date, no such experiment has been performed *in vivo*, but many other studies have used axon activity as a proxy for ACh levels^15,16,18,20,22^. Broadly, we confirmed that local ACh levels do indeed correspond to axon activity as expected, and that the longer-lasting increases in ACh seen during running are much slower to decay than the rapid, phasic increases linked to dilation. This suggests that, in line with pharmacological studies indicating the rapid action of acetylcholinesterase (AChE)^35^, small amounts of ACh are cleared quickly but larger increases may transiently saturate clearance mechanisms leading to longer decays^36^. Consistent with this scenario, we found that the timing and magnitude of small increases of ACh around pupil dilation vary depending on dilation magnitude.

Spatially, we found that ACh availability decreases with increasing distance from axons, consistent with volume transmission of ACh. There has been an ongoing debate about whether ACh can escape the synapse and influence receptors on neighboring processes not directly synapsing onto a cholinergic axon^36,37^. Our finding that the magnitude of cholinergic spike-triggered ACh activity displays a distance-dependent relationship is consistent with the idea that these experiments represent the first direct observation of volume transmission in vivo. Overall, our simultaneous GRAB-ACh/axon recordings support the notion that cholinergic axon activity can indeed be used as an accurate proxy for changes in local ACh levels, which is reassuring given the many existing studies that have used ChAT-expressing axons as a proxy for ACh.

Given the close correspondence of ACh activity with axon activity and pupil size or locomotion, we wanted to quantify the extent to which it is possible to predict ACh levels from these variables. Predictive modelling revealed that cholinergic axon activity and behavioral state variables, especially pupil size, can be used for accurate predictions of cortical ACh levels, explaining the majority of variance on held-out data. Pupil and treadmill activity is routinely recorded in many head-fixed mouse experiments, and so this model may be useful for inferring ACh fluctuations in the absence of ground truth training data.

Throughout this study, we uncovered a close link between pupil size and cortical ACh levels. This was somewhat surprising, as traditionally the locus coeruleus (LC) noradrenergic system is thought to be the neuromodulator most closely linked to pupil size^27^. The anatomical pathway by which LC could control pupil size has been well-established, but it is less clear by what pathway pupil size could be linked to ACh. It’s possible that fluctuations in cortical ACh delivered via the BF may occur downstream of projections from LC to BF^38,39^. Further studies are needed to examine this possibility, but we hope that this study emphasizes the translational potential of pupillometry for inferring fast changes in ACh, and that caution should be taken when using fluctuations in pupil size as a proxy for exclusively noradrenergic activity in humans.

Our study provides a bridge between previous work using cholinergic axonal GCaMP imaging and recently developed genetically encoded fluorescent sensors, and provides additional validation of GRAB-ACh for investigating the question of when and where ACh is available in the brain. The deconvolution approach we demonstrate here may be useful for other situations where it is desirable to estimate the time course of changes in levels of a ligand while removing the temporal smoothing due to slow sensor kinetics. Overall, these experiments contribute to a growing understanding of the spatiotemporal scale of ACh availability in the cortex in relation to state-related behavioral variables. This understanding is essential for future work focused on studying the downstream effects of ACh in changes on cortical excitability and information processing during perception and behavior.

## METHODS

### Animals, surgery, and injection of viral vectors

All procedures were carried out in accordance with the ethical guidelines of the National Institutes of Health and were approved by the Institutional Animal Care and Use Committee (IACUC) of Baylor College of Medicine. This study used a total of 32 mice aged 1.5 to 8 months (n=14 male, n=18 female).

#### Surgical procedures for spontaneous-state-change experiments and ACh injection experiments

Wild type C57Bl/6 mice (or various other negative genotypes on a C57Bl/6 background) were implanted with a 4mm cranial window over visual cortex (n=16), a 5mm window spanning visual and somatosensory cortices (n=3), or a 3mm window over motor cortex (n=2) under isoflurane anesthesia. The head was shaved over the dorsal portion of the skull and bupivacaine 0.25% (0.1 ml) and ketoprofen (5 mg/kg) were administered subcutaneously. The skin was pulled back and glued in place using Vetbond (3M). A headbar (steel washer with outer diameter 0.438”, inner diameter 0.314”, thickness 0.016”) was affixed to the skull using dental cement (C&B Metabond). A micromotor rotary drill (Foredom) fitted with a 0.6mm tungsten carbide bur was used to thin the skull over the center of the chosen cortical area, which was determined by stereotactic coordinates (VIS: 2.8 mm lateral, 1mm anterior lambda; SS: 2.8mm lateral, 2.25mm posterior bregma; MO: 1.5mm lateral, 1mm anterior bregma), and a circle of skull was removed. A borosilicate glass pipette (World Precision Glass) was pulled to a tip width of 40µm and attached to a syringe which was used to inject 200-2000nl of AAV-hSyn-ACh4.3 at a depth of 200-700µm below the surface using a Micro4 MicroSyringe pump controller. The dura was removed, and a #1 glass coverslip (thickness 0.15 ± 0.02 mm) was then glued in place using super glue (Loctite). The animal was allowed to recover in its home cage on a heated disk until sternal recumbency was regained, and was monitored for the next 3 days for signs of pain or distress.

#### Surgical procedures for dual axon-ACh imaging experiments

ChAT-cre (n=1) or Δneo ChAT-cre (n=4) mice first received an injection into the horizontal diagonal band (HDB) of the basal forebrain after anesthesia induction by i.p. Injection of ketamine and xylazine. HDB was selected due to its preferential cholinergic projections to the visual cortex^24,29,30^. The head was shaved and skin pulled back after an incision was made. A burr hole was drilled and, micropipette fitted with a 20µm glass pipette was positioned to AP: 0.13, ML: 1.1, DV: -5.6, and 1600nl of AAV-syn-FLEX-jGCaMP8s was injected over 160 seconds. The skin was sutured back in place using braided absorbable sutures (McKesson) and the animal allowed to recover. After 1 week, a headbar and 4mm cranial window was implanted over the visual cortex using the above procedures, into which 1000-1800nl of AAV-hSyn-rACh1.0 was injected at a depth of 300µm below the surface.

### In vivo pharmacological experiments

In 4 mice under general isoflurane anesthesia, the glass coverslip previously implanted over the visual cortex was removed and replaced with a coverslip of the same size but with a small hole near the edge. The mouse was head-fixed under the microscope where it remained anesthetized for the duration of the experiment. The following agents were diluted with cortex buffer and Alexa 568 dye to achieve specific molecular concentrations: acetylcholine chloride (crystals/powder, Sigma-Aldrich), norepinephrine bitartrate (crystalline, Sigma-Aldrich). A borosilicate patching pipette was fitted to a syringe and backfilled with the mixed pharmacological agent and inserted into the cortex through the hole in the coverslip. 200 mBar pressure was applied to the syringe, which was released with a button press.

### Histology

After the completion of two-photon imaging, mice were deeply anesthetized with isoflurane and transcardially perfused with 4% paraformaldehyde (PFA). Brains were removed from skulls and stayed in 4% PFA overnight at 4°C. 100-µm-thick coronal sections were cut with a vibratome (Leica 1000S) in 0.1 M PBS. Unspecific antigens were blocked by incubating selected sections for 1 hour in 10% normal donkey serum (NDS, Sigma, Cat# D9663) and 0.5% Triton-X-100 in 0.1 M PBS. Sections were selected that contained the HDB injection sites and Flex-jGCaMP8s labeled axons in VIS. Following the blocking step, the selected sections were incubated in a 1:300 dilution of goat polyclonal antibody against ChAT (Millipore Cat# AB144P, RRID: AB_2079751), 2% NDS and 0.5% Triton-X-100 in 0.1 M PBS overnight at 4°C. On the second day, the sections were washed with 0.01 M PBS three times for 10 mins each and then incubated in a 1:100 dilution of a rabbit-anti-goat antibody that was directly conjugated to AlexaFluor-546 (Invitrogen, Cat# A-21085, RRID: AB_2535742), 2% NDS and 0.5% Triton-X-100 in 0.1 M PBS for 2 hours. The sections were then washed with 0.01 M PBS and mounted on slides using VECTASHIELD antifade mounting medium containing DAPI (VectorLabs, Cat# H-1200-10).

### Locomotion and pupillometry

For all experiments, the mouse was positioned atop a 9”-diameter x 3”-wide cylindrical styrofoam treadmill and head-fixed using a steel headbar clamped onto the surgically implanted washer. The surface of the treadmill was covered in duct tape to provide extra traction. The mouse’s body was free to move with 2 degrees of freedom (forward and backward). An optical encoder (McMaster-Carr) was used to keep track of treadmill speed. Treadmill traces were processed with a 0.5 second median filter. Running periods were defined as running at a speed of least 1cm/sec for at least 1 second. Periods less than 3 seconds apart were grouped together into a single period. Periods of quiescence were determined by removing all running periods - 3 seconds.

A Dalsa camera mounted with a 55mm telecentric lens (OEM) and a 1” hot mirror (Thorlabs) was used to capture a video of the mouse’s eye at 20fps. As the experiment was conducted in darkness with no visual stimulus, the eye was illuminated from an outside source (either a computer monitor at or a UV light) to prevent total pupil dilation and allow normal, spontaneous fluctuations in pupil size to be measured.

A DeepLabCut model was used to identify pupil edges from the eye videos, from which the circumference of the pupil was interpolated. The radius of the pupil (in pixels) was then determined, and this trace was 1hz lowpass filtered. Radius traces were inspected by eye to determine quality of tracking, and movies which had poor tracking were thrown out. To convert radius in pixels to radius in millimeters, the corners of the eyelid were detected using DeepLabCut and the median distance was assumed to be 2.757mm (which was determined by our measurements to be the universal mouse eyelid width). This was used to create a conversion factor whereby pixels could be converted to millimeters. Dilation (or constriction) events where then defined as periods of increasing (decreasing) pupil radius lasting greater than one second and with an average speed greater than 0.02mm/sec. Blinks or frames where the pupil was not able to be detected by DeepLabCut were considered NaNs, and any event comprised of more than 15% NaNs was dropped.

### Imaging

All scans were recorded using a two-photon fast resonant scanning system (ThorLabs). Frame rates varied from 5-60fps. Excitation wavelengths were either 920nm (GRAB-ACh scans) or 1000nm (GCaMP and rACh scans) with a 25x objective (Nikon). Output power was kept around 15-40mW.

### Motion correction and selection of ROIs

All scans were corrected for motion in the X and Y directions via a two-step algorithm. Large scale motion was calculated by taking the average of the movie and computing phase correlation between this template and each frame of the movie. After correcting for large scale motion, smaller scale motion was corrected by taking the average of a 2,000-frame window and phase correlating frames within this window to the local template. Windows were selected to have 500 overlapping frames, and a linearly weighted average was taken for the displacement in the overlapping region.

Scans were discarded which had poor motion correction (defined as root-mean-squared of the x- and y-shifts produced from motion correction above 4 microns) or which contained large, unexplained jumps in fluorescence (at least 20x the average derivative). Three scans were discarded in which, despite adequate motion correction, it was assumed that the recorded axons moved out of the recording plane during motion due to a complete absence of detected spikes during running periods.

Fluorescence traces for each scan were determined by taking the average fluorescence of all pixels within a certain ROI (“mask”). For scans which recorded GRAB-ACh or rACh sensors, masks were determined by taking pixels which had average brightness values <99.9^th^ percentile and >(99.9^th^ percentile/2) or >(99^th^ percentile/2), respectively, after scan borders were removed (see Supp. Fig. 3). The more inclusive lower-bound threshold used for the rACh scans was selected due to overall dimmer expression of the red sensor versus the green sensor. Each mask was then eroded using a 7x7-pixel kernel to provide smoothness. For scans which contained axons, a single mask was determined for the green (GCaMP) channel by calculating the overlap between a manually drawn mask over visible axons and a brightness mask (i.e. pixels with an average brightness value greater than (99^th^ percentile/2) after removing borders). For scans where ACh was injected, a circular mask with 50µm radius was created over the tip of the injection pipette, which was manually defined.

For distance-binned peri-spike traces, six masks for the red (rACh) channel were determined for each scan (see Fig. 5A). First, the distance in µm from each pixel to the nearest pixel which contained an axon (as defined by the axon mask described above) was calculated. Five distance bins were determined based on the distribution of distances across all scans, excluding axon-containing pixels (see Supp. Fig. 1). Then masks were calculated by sorting all pixels in each scan into one of six bins (axon-containing pixels/”bin 0”, or one of the five distance bins).

### Characterization of and control for hemodynamic artifact

Hemodynamic artifacts are always a concern when recording bulk fluorescence with a sensor such as GRAB-ACh (although much less of a concern than with one-photon methods). With GRAB-ACh, we have found such an artifact in the form of a brief decrease (“dip”) in fluorescence which specifically occurs at the onset of locomotion. This dip is observed both with non-responding mutated versions of the sensor and also with GFP. In our characterization of this artifact, we’ve discovered that this dip is more prominent when recording at deeper cortical depths than at superficial depths (Supp. Fig. 4A & 4B), as expected for fluorescence signals collected through larger (deeper) volumes of tissue. To be conservative, we excluded GRAB-ACh scans from our analysis which were recorded at or deeper than 100µm in VIS (a small number of scans recorded between 100-200µm in SS and MO were left in the analysis due to the limited number of scans in those areas).

To further characterize this artifact, we performed both 2-channel and 2-field (i.e., 2-ROI) controls with GRAB-ACh-mut and rACh or GRAB-ACh (Supp. Fig. 4C & E). In the 2-channel recordings, although a run-related artifact was found in GRAB-ACh-mut, no such dip was found in the rACh signal (Supp. Fig. 4D). In this case, correcting the rACh trace with the mut trace did not result in qualitative or quantitative differences in the rACh signal. In the two-field recordings, a run-related artifact was observed in both fields (Supp. Fig. 4F. While correcting the GRAB-ACh trace with the mut trace did remove the artifact from run onset, it again did not result in qualitative or quantitative differences in the GRAB-ACh signal.

Both of the above correction methods are potentially flawed. When imaging two simultaneous fields, the differing vasculature patterns between the two sites could potentially lead to differences in artifact characteristics leading to an incorrect correction. When imaging two channels, the different fluorophores (red vs. green) could similarly contribute to differences in artifact characteristics. After extensive characterization, we believe that for the GRAB-ACh sensor, the signal to noise appears to be relatively high - we did not observe a prominent hemodynamic artifact with rACh, and the two-field correction with GRAB-ACh-mut did not substantially alter the characteristics of the GRAB-ACh signal. Thus, our approach is to present the minimally processed fluorescence after removing deeper scans that might be more impacted by hemodynamic changes. However, caution should be used with less sensitive sensors with signals that may be dominated by vascular artifacts.

### Peri-dilation, peri-run, and peri-spike traces

Fluorescence traces were first preprocessed as follows. The first 20 seconds of each trace was thrown out in order to account for stabilization of the laser. Traces were then detrended to correct for photobleaching by subtracting a 4^th^-order polynomial, 1hz lowpass filtered (except for Fig. 5B, which used a 2hz lowpass filter), and converted to ΔF/F. Baseline F was calculated using a 20-minute non-causal rolling 5^th^-percentile and ΔF/F was calculated as (fluorescence - baseline)/baseline. Traces which had negative baseline values were discarded in order to avoid dividing by a negative baseline. Deconvolved traces in Fig. 6 were preprocessed in this order: detrending, deconvolution (see method in section below), filtering, ΔF/F normalization.

The times at which dilation, run onset, and run offset events occurred were determined using the methods in the above section (“Locomotion and pupillometry”). Peri-event traces were calculated by subtracting the average pre-event fluorescence value (“baseline”) and multiplying by 100 to convert to a percentage. Peri-dilation events containing NaNs in the radius trace were excluded. All peri-event traces were averaged together and standard error was found by dividing the standard deviation of all traces by the square root of the number of traces.

To determine spike times for peri-spike traces, the raw fluorescence calculated from the axon mask described above (“Motion correction and selection of ROIs”) was deconvolved using a non-negative matrix factorization algorithm^40^. For each trace, a threshold was set at the 96^th^ percentile of the deconvolved fluorescence values and every sample point at or above this threshold was defined as a spike time. This threshold level was selected in order to achieve spiking rates which matched biologically-plausible firing rates of the cholinergic basal forebrain (approximately 4 spikes/sec overall^41^ and approximately 7 spikes/sec during active periods^42^).

### Coherence and cross-correlation

Coherence between pupil radius and ACh sensor activity or GCaMP axon activity was performed on unfiltered radius and fluorescence traces using a 1-minute (1,200 sample) long Discrete Prolate Spheroidal Sequences window with overlap of 1,100 and half-bandwidth of 4. Fluorescence traces were detrended and radius traces had NaNs interpolated before calculating coherence.

Cross-correlations were performed by first normalizing pupil radius and fluorescence traces. Unfiltered and NaN-interpolated pupil radius traces were normalized by subtracting the mean and dividing by the standard deviation times the number of samples in the trace. Unfiltered and detrended fluorescence traces were normalized by subtracting the mean and dividing by the standard deviation of the trace. Cross-correlations were then calculated on the normalized traces, and peaks/lags were determined using the middle +/- 2 seconds of the cross-correlations.

### GRAB-ACh deconvolution

#### Fluorophore impulse response

Recent empirical work had characterized the GRAB-ACh sensor kinetics^23^; first used those results to model the fluorophore’s impulse response (or convolution/deconvolution kernel) *k*. The exponential fall time had been directly determined to be 580*ms*, by measurement of fluorescence in the presence of a competitive agonist. The inverse rise-time of the fluorescence response to an ACh puff increases logarithmically with molarity, with a constant 3.12/(*μM* × *s*). However, as the convolution kernel is defined as the response to an instantaneous pulse of infinite concentration, and moreover because these are association kinetics (rather than kinetics of conformation state change in the fluorophore itself) by extrapolation the kernel’s rise time should be effectively zero; we therefore modeled the kernel as a single falling exponential with time constant 580*ms*.

#### Convolutional model

We then assumed (cf. Yaksi and Friedrich, 2006^43^) that the fluorescence signal is generated by convolution of a time-varying concentration [*ACh*](*t*) with the fluorophore’s impulse response *k*, so that the GRAB-ACh fluorescence trace *F*(*t*) is:

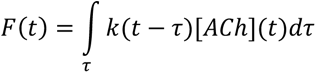

Roughly, the instantaneous concentration [*ACh*] at time *τ* will cause a characteristic fluorescence response *k*(*t* − *τ*) --- which lasts for some time after fluorophores are activated. Contributions from fluorophores so activated at various times in the past (i.e., *t* − *τ*) are then integrated to generate the fluorescence at time *t*. Note that this model does not account for nonlinear effects, in particular saturation.

#### Deconvolution

Deconvolution methods then attempt to invert the process of convolution, recovering the source [*ACh*] from the fluorescence signal and the impulse response of the fluorophore. The simplest deconvolution method divides the Fourier transform (denoted ℱ, with inverse transform ℱ^$(^) of the fluorescence signal by that of the kernel:

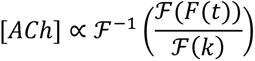

However, we found this approach led to estimates of [*ACh*] which – at least at the level of individual scans -- were so noisy as to be uninterpretable.

More advanced deconvolution methods differ largely by their assumptions on source and noise, yielding a denoised estimate of a conditioned source. Several modern methods (see also the review^44^), for instance, deconvolve latent voltage spikes from calcium fluorescence^45^. In our study, however, we cannot assume *a priori* that acetylcholine concentration [*ACh*] is sparse or spikelike. A classical approach --- Wiener deconvolution (which had also previously been applied, with some modification, in the calcium context^46^) --- yielded intelligibly denoised single-trial [*ACh*] estimates without further source constraints, making it a suitable method for our purposes.

Wiener deconvolution modifies the division method above by adding a regularization parameter to the denominator:

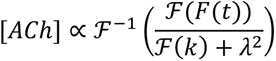

We estimated this parameter *λ* based on empirical considerations: As it increases, the reconstructed signal first shows removal of higher-frequency noise, but then shows a flattening of the variation of interest at lower frequencies --- eventually approaching a constant value (See Supplemental video 3). To select an appropriate degree of smoothing for each individual scan, we first standardized the signal, then varied lambda until the absolute reconstruction error was 10% of its maximum (i.e. 10% of when the signal flattened completely). Although this ‘optimal’ value of lambda was slightly larger computed over entire scans vs. stationary-only periods (*p* < 10^−10^; Wilcoxon test), the size of this difference was fairly small (mean: 11.3%).

### Axonal to ACh predictive modelling

The axonal to ACh predictive model related the axonal fluorescence (*F*) to the simultaneously recorded rACh signal. First, a sigmoidal function calculated the maximal axonal output, *S*(*F*),

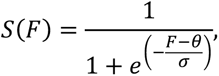

where *θ* horizontally shifts the sigmoid (e.g., high *θ* → smaller output *S* for given activity *F*) and *σ* is the rate of growth of the sigmoid from baseline to threshold. Next, we simulated the temporal release of ACh following an exponential rise and decay function by temporally convolving *S*(*F*) with the release function *R*(*t*),

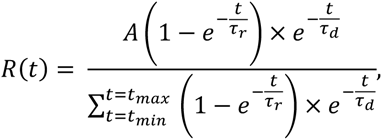

where *τ_r_* and *τ_d_* are the release rise and decay times, *A* is a scaling parameter, and the denominator is a normalization factor. Finally, the predicted ACh is convolved with the sensor dynamics 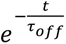 (normalized to unit area) with *τ_off_* = 245 ms (estimated from unpublished results obtained from personal communications). The optimization swept across the 5 parameters (*θ*, *σ*, *A*, *τ_r_*, *τ_d_*) using an interior-point algorithm (MATLAB fmincon) over 150 iterations to minimize the RMSE between the predicted and empirically recorded GRAB-ACh signal.

### Behavioral to ACh predictive modelling

We predicted GRAB-ACh fluorescence from behavioral signals – running speed, pupil size and temporal derivatives – using a long short-term memory (LSTM) recurrent neural network. Briefly, the RNN consisted of an input layer, an LSTM layer (with a working memory of the previous ∼ 8.5 s), a fully connected 256-node recurrent neural network, a 10% dropout layer, and a fully connected output layer. The model was trained on ∼ 2-minute epochs across 90% of the recordings and validated on unseen held-out recordings (10% of total recordings). We optimized the parameters of the network through stochastic gradient descent using the Adam optimizer over 150 iterations, minimizing the RMSE between predicted and observed GRAB-ACh. The predictive model was trained using running speed, pupil size and their temporal derivatives (pupil + speed), pupil size and its temporal derivative (pupil), and running speed and its temporal derivative (speed). Model fits were assessed by calculating the Pearson correlation between predicted and observed GRAB-ACh in the held-out validation dataset.

## ACKNOWLEDGEMENTS

We would like to thank the Andreas Tolias lab for providing resources and equipment. E.N. was supported by NIH NRSA Fellowship F31NS122428. C.S. was supported by NIH NRSA Fellowship F31NS132465. Y.L. was supported by The National Science Fund for Distinguished Young Scholars and NIH BRAIN Initiative (1U01NS103558, 1U01NS113358, and 1U01NS120824). M.J.M was supported by NIDCD awards R01 DC017797 and R03 DC015618. J.M.S was supported by the National Health and Medical Research Council (GNT1193857) and the Bellberry Foundation. J.R. was supported by NIH awards 1R34NS132045 and 1RF1NS128901.

## AUTHOR CONTRIBUTIONS

E.N. performed experiments, analyzed data, and wrote the paper. N.Z. performed subcortical injections and histology. B.R.M. carried out the predictive modelling. R.G.L. developed the sensor deconvolution method. C.S. helped develop and troubleshoot the sensor preprocessing pipeline. Z.H.M. conducted experiments. F.A.B. performed anatomical imaging. G.L. assisted with sensor development. Y.L. supervised sensor development and provided resources. M.J.M. and J.M.S. provided supervision. J.R. designed the study, performed experiments, and provided funding acquisition, supervision, and review/editing.

---

**Supplemental Fig. 1.**
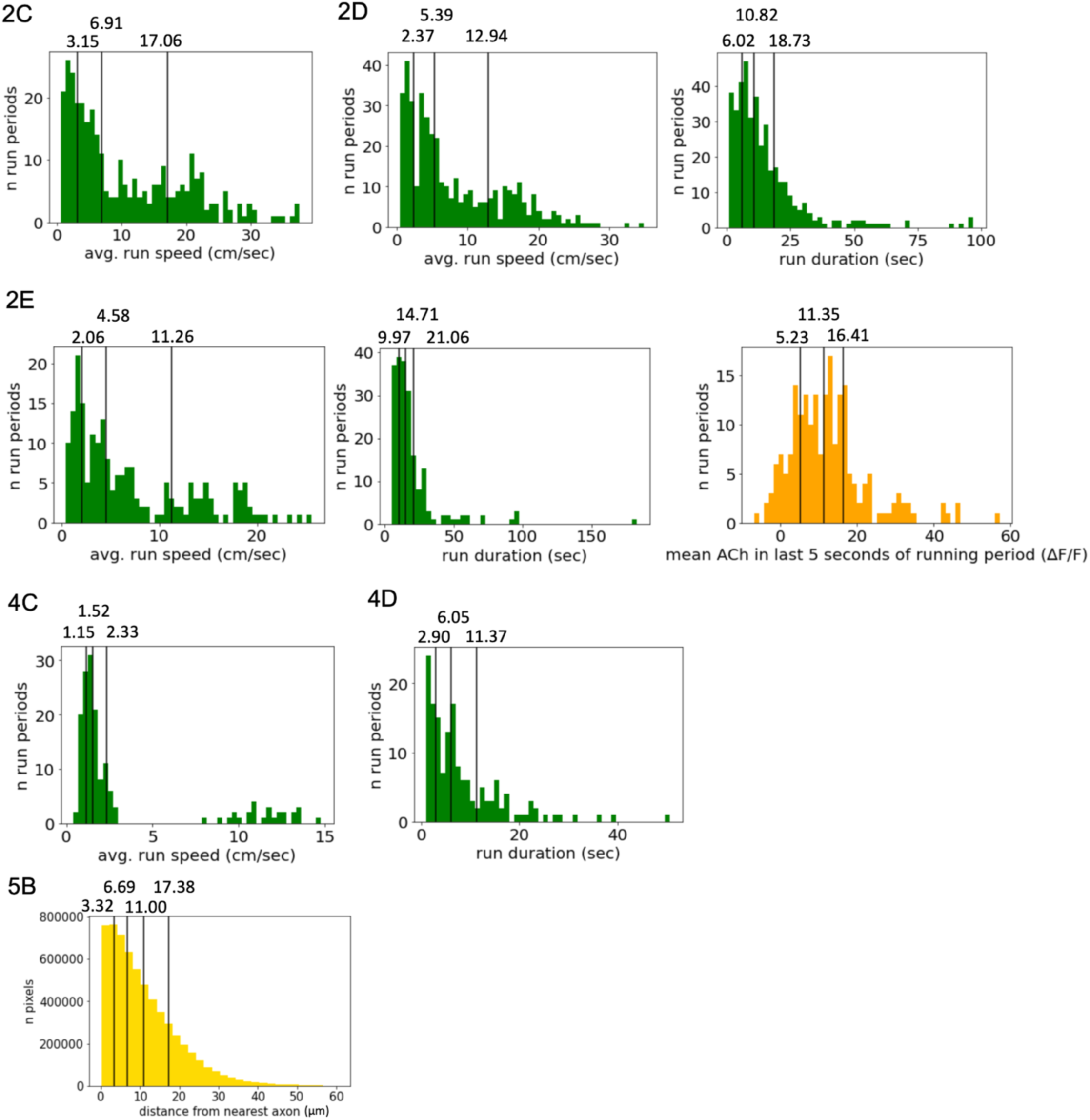
Run qualities, peak ΔF/F, and distances per quartile/quintile for. Figures 2**, 4 & 5.**

**Supplemental Fig. 2.**
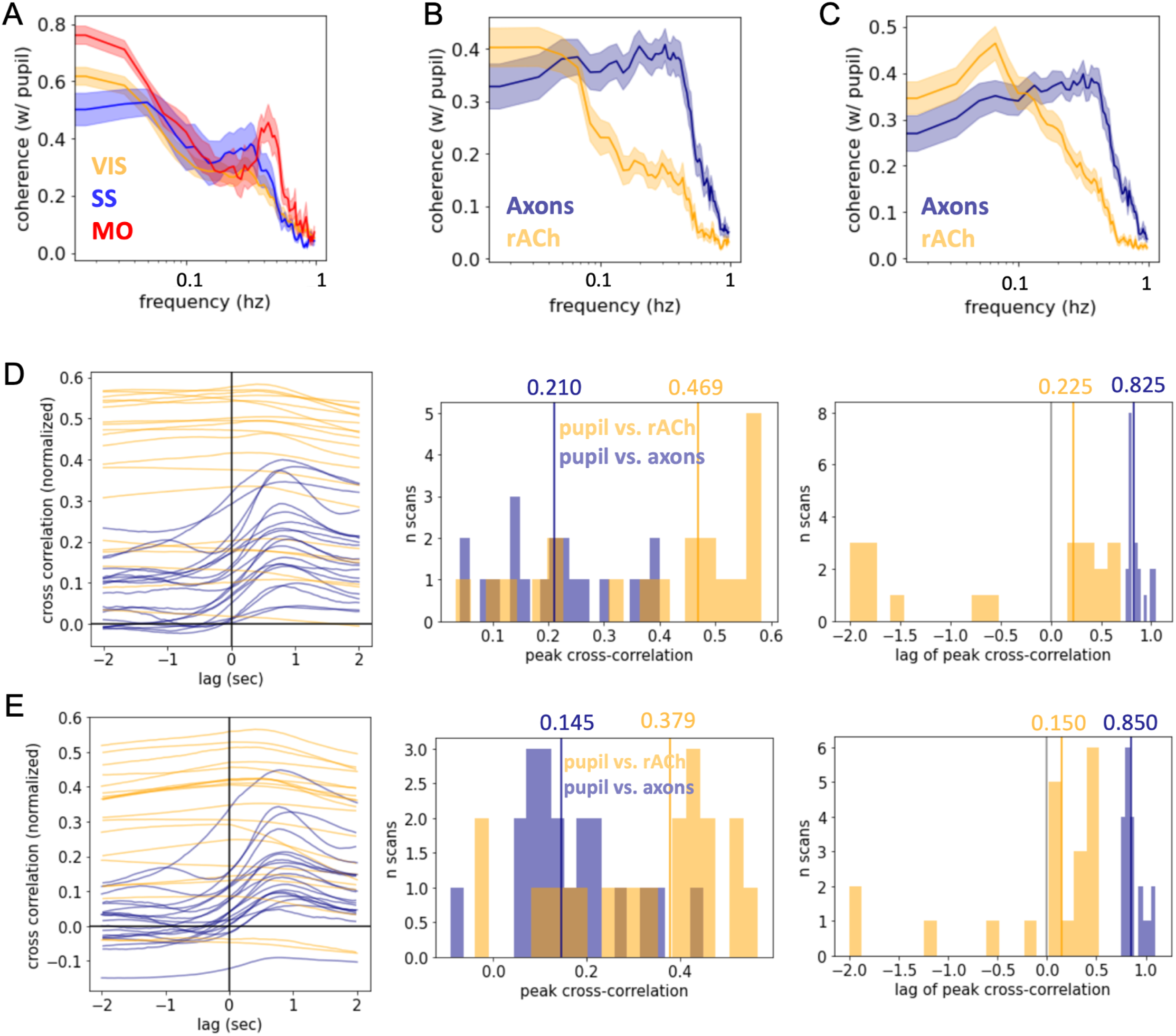
Coherence and cross-correlation of pupil with ACh or axon activity. **A)** Mean coherence of pupil radius with ACh activity in VIS, SS, and MO during full scans (i.e., including running periods). **B)** Mean coherence of pupil radius with cholinergic axon and ACh activity during full scans. **C)** Mean coherence of pupil radius with cholinergic axon and ACh activity during periods of quiescence. **D)** Cross-correlations for full scans. (left to right: cross-correlation traces for each scan; distribution of peak cross-correlations with median values above; distribution of lags with median values above). **E)** Cross-correlations for quiescent periods (left to right: as above).

**Supplemental Fig. 3.**
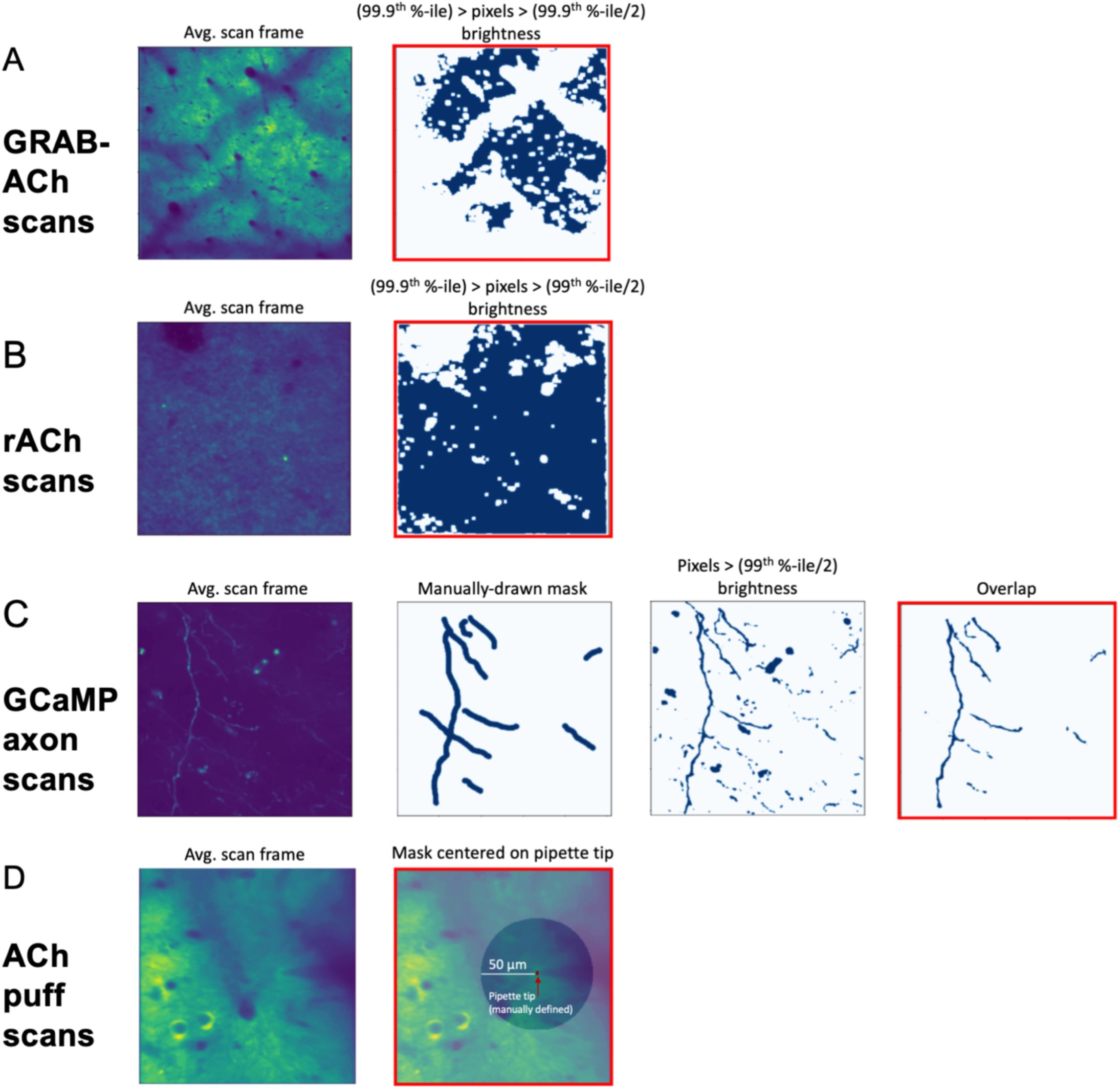
Determination of ROIs. In each panel, the red-boxed image was the final ROI used. **A)** Example average scan image and resulting mask for a GRAB-ACh scan. **B)** Example average scan image and resulting mask for an rACh scan. **C)** Example average scan image, the mask manually drawn over axons, the brightness mask, and the overlap of the two. **D)** Example average scan image and mask resulting from taking a 50µm radius around the manually defined pipette tip.

**Supplemental Fig. 4.**
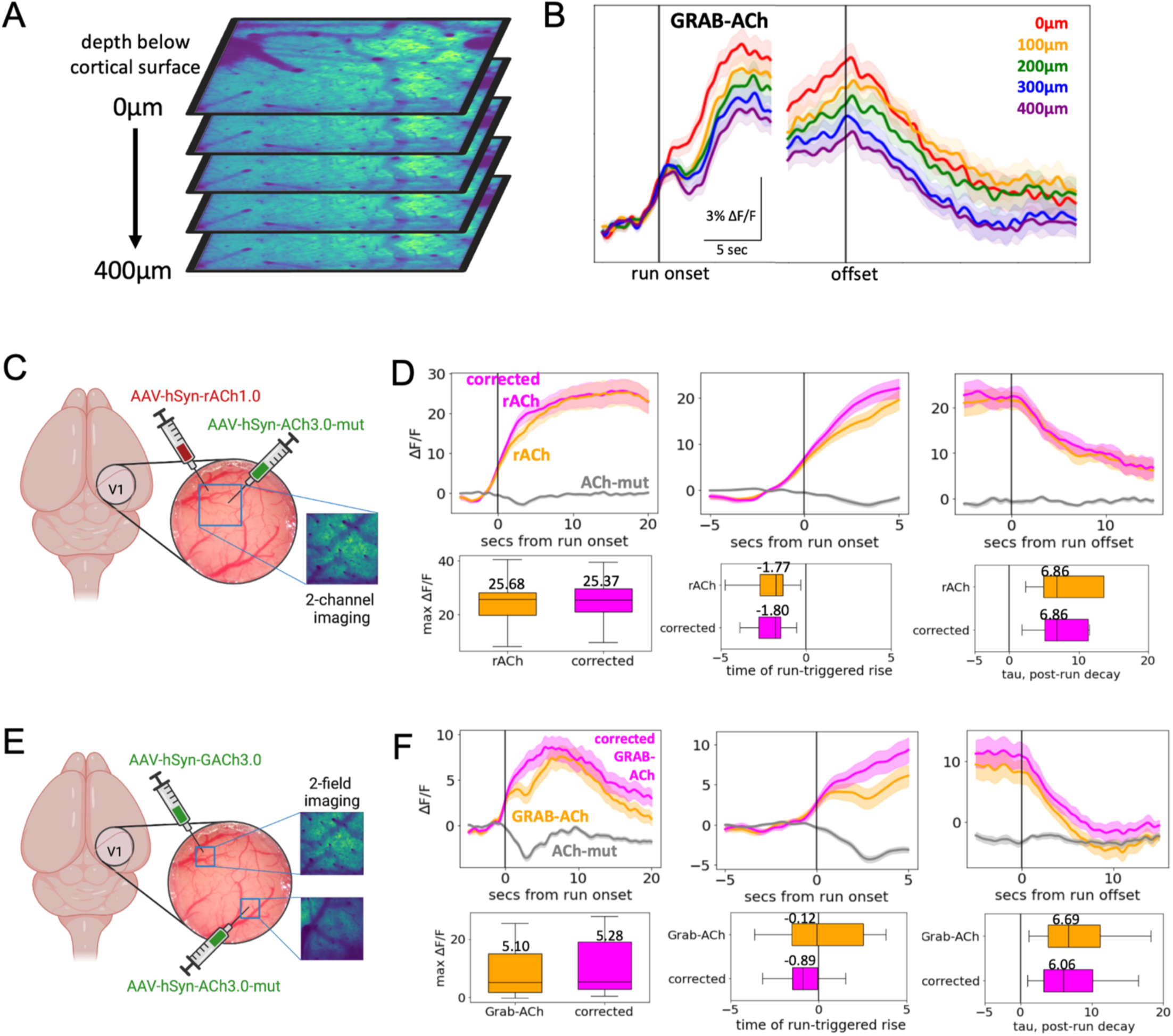
A locomotion-linked hemodynamic artifact found in the GRAB-ACh sensor is more prominent at deeper depths and does not substantially affect measured ACh dynamics. **A)** Certain GRAB-ACh control scans were collected with 5 simultaneous planes in the Z direction. **B)** Run onset- and offset-triggered GRAB-ACh fluorescence at 5 simultaneous depths. **C)** A two-channel imaging method to control for hemodynamic artifacts with rACh. rACh and GRAB-ACh-mut were expressed in the same field and imaged simultaneously on different channels. The “corrected” trace was computed by adjusting the mean of the GRAB-ACh-mut trace to match the mean of the rACh trace (outside of running periods), then subtracting this adjusted mut trace from the GRAB-ACh trace. **D)** Run onset- and offset-triggered rACh and corrected rACh with corresponding maximum response, rise onset time, and decay tau. All pairwise comparisons of medians were not significantly different (Kruskal-Wallis test). **E)** A two-field imaging method to control for hemodynamic artifacts with GRAB-ACh. GRAB-ACh and GRAB-ACh-mut were expressed in spatially segregated areas of the same window and imaged simultaneously in different ROIs. The “corrected” trace was computed as in panel D). **F)** Run onset- and offset-triggered GRAB-ACh and corrected GRAB-ACh with corresponding maximum response, rise onset time, and decay tau. All pairwise comparisons of medians were not significantly different. Panels c and e created with BioRender.com

